# *Dnmt1* has an essential function despite the absence of CpG DNA methylation in the red flour beetle *Tribolium castaneum*

**DOI:** 10.1101/426890

**Authors:** Nora KE Schulz, C Isabel Wagner, Julia Ebeling, Günter Raddatz, Maike F Diddens-de Buhr, Frank Lyko, Joachim Kurtz

## Abstract

Epigenetic mechanisms, such as CpG DNA methylation enable phenotypic plasticity and rapid adaptation to changing environments. CpG DNA methylation is established by DNA methyltransferases (DNMTs), which are well conserved across vertebrates and invertebrates. There are insects with functional DNA methylation despite lacking a complete set of *Dnmts.* But at least one of the enzymes, DNMT1, appears to be required to maintain an active DNA methylation system. The red flour beetle, *Tribolium castaneum,* lacks *Dnmt3* but possesses *Dnmt1* and it has been controversial whether it has a functional DNA methylation system.

Using whole genome bisulfite sequencing, we did not find any defined patterns of CpG DNA methylation in embryos. Nevertheless, we found *Dnmt1* expressed throughout the entire life cycle of the beetle, with mRNA transcripts significantly more abundant in eggs and ovaries. A maternal knockdown of *Dnmt1* caused a developmental arrest in offspring embryos.

We show that *Dnmt1* plays an essential role in *T. castaneum* embryos and that its downregulation leads to an early developmental arrest. This function appears to be unrelated to DNA methylation, since we did not find any evidence for this modification. This strongly suggests an alternative role of this protein.

## 1 Introduction

Phenotypic plasticity is enabled by epigenetic mechanisms, which help the organism to adapt rapidly to changing environmental conditions without affecting the underlying genotype ^1^. One of these mechanisms is DNA methylation, which in insects is associated with a wide variety of phenotypically plastic traits ^2,3^. In eusocial insects, caste formation can correlate with differential DNA methylation patterns ^4–6^. One of the first examples for functional DNA methylation in insects was the discovery that the downregulation of the methylation machinery in honey bee larvae caused them to develop into a queen-like phenotype, which is naturally established by the feeding on royal jelly ^7^. Furthermore, DNA methylation patterns do not only differ between the brains of queens and workers ^5^, but also within the worker caste, where they have been associated with different tasks performed by the worker bee^8^. Additionally, these epigenetic marks appear to be reversible if the individual has to switch its task ^9^. Furthermore, in two ant species the same set of genes was differentially methylated according to developmental stage and caste ^6^. DNA methylation can also be observed in non-social insects and may contribute to the striking phase polyphenism between solitary and gregariously living migratory locusts ^10,11^. On the other hand, it also has been suggested that DNA methylation can have a more general, basic role in insects and it might function in more ubiquitously essential processes, e.g. embryonic development, genomic imprinting and alternative splicing ^6,12–15^.

Insect methylation patterns can be highly diverse ^15,16^. In cockroaches and some lepidopterans, DNA methylation can reach levels similar to those observed in vertebrates and plants ^15,17^. This is also true for some regions of the genome of the stick insect, but here in addition to gene bodies also repetitive elements are strongly methylated ^18^. In contrast, holometabolous insects possess sparser DNA methylation than vertebrates ^15,16,19^. Most of these insects also differ from mammals in the location of their methylation marks, as the majority of CpG methylation occurs within gene bodies ^6,20,21^. The placement of methyl groups at these mostly intragenic loci could hint at its possible role in alternative splicing ^22^.

In invertebrates and vertebrates alike, the same conserved set of enzymes adds methyl groups to cytosines followed by a guanine (CpG) ^23^. The resulting CpG DNA methylation patterns are therefore symmetrical on both strands of the DNA. In most cases, two DNA methyltransferases (DNMTs) are needed to sustain functional CpG DNA methylation. DNA methyltransferase 1 (DNMT1) is responsible for maintaining pre-existing methylation patterns by methylating hemimethylated DNA after replication, while DNA methyltransferase 3 (DNMT3) establishes new methylation patterns on previously unmethylated cytosines ^23^. Previously, it was thought that a third enzyme, DNMT2, was also involved in DNA methylation because of sequence conservation and the fact that it also contains a cytosine methylase domain ^24^. However, several studies have shown that this enzyme methylates small RNAs, especially tRNA ^24,25^. Furthermore, organisms that lack *Dnmt1* and *3,* but contain *Dnmt2* do not possess a functional DNA methylation system ^24,26,27^.

The distribution of *Dnmts* in insects is quite diverse. While many insects, especially eusocial species, have a complete set of *Dnmts,* some with an expansion in copies for *Dnmt1,* other insects completely or partially lack *Dnmts* and some also fail to produce functional levels of DNA methylation ^15,19,28^. Diptera appear to have lost both *Dnmt1* and *Dnmt3* and with it any relevant CpG DNA methylation ^15,16^. Some insect species including examples from Coleoptera, Hemiptera, Lepidoptera and even social Hymenoptera are lacking a gene encoding for DNMT3 ^15,29–31^. Nevertheless, several of them may show levels of DNA methylation, which presented either directly in sequencing studies or were deduced from the CpG depletion in their genomes ^15,16,30^. Thus, the question arises, how a functional DNA methylation system can be sustained in the absence of an enzyme able of *de novo* methylation. It has been suggested that DNMT1 could take over this function, but to the best of our knowledge there are no studies demonstrating this.

The status of a functional DNA methylation system in the red flour beetle *Tribolium castaneum* has been controversially discussed over the past years. Over evolutionary time, the mutagenic effect of a methyl group attached to a cytosine leads to a depletion of CpGs from the genome ^32^. However, the beetles’ genes do not exhibit the bimodal distribution of normalized CpG content associated with the presence of DNA methylation in other insects ^4,15^. Additionally, in adult beetles, low-coverage whole-genome bisulfite sequencing (WGBS), which reveals methylation at single-base resolution, could not detect any defined patterns of CpG DNA methylation ^19^. On the other hand, using an approach based on methylation sensitive restriction endonucleases, which lacks the sensitivity and specificity of WGBS, Feliciello *et al.* (2013) found evidence for DNA methylation in early embryos, whereas larvae, pupae and adults were found to be mostly devoid of methylation. While this raised the possibility of an active demethylation process in later developmental stages, resembling the epigenetic reprogramming in mammalian embryos ^34,35^, additional bisulfite sequencing of satellite DNA indicated that these fragments remain heavily methylated throughout the entire lifetime of the individual, even in non-CpG contexts ^33^. Furthermore, another study based on bisulfite sequencing also found DNA methylation outside the context of CpG and mostly at CpA ^36^.

The controversial evidence of CpG DNA methylation in *T. castaneum* demands for additional, functional studies in this important insect model organism. In *T. castaneum* we observe phenotypic plasticity in the form of immune priming, *i.e.* a more successful response to a pathogen at a secondary exposure ^37^. This immune priming can be maternally and paternally transferred to the offspring ^38–40^, which makes the involvement of epigenetic modifications in this phenomenon likely. Besides other epigenetic mechanisms, DNA methylation and its facilitating enzymes have been indicated to play a role in the evolution of resistance and the rapid adaption we see during host-parasite co-evolution ^41,42^. Therefore, we here investigate the existence of CpG DNA methylation and the role of *Dnmt1* in *T. castaneum* using WGBS and knockdown of *Dnmt1* using maternal RNAi.

## 2 Results

### 2.1 WGBS confirms the absence of CpG methylation in *Tribolium* eggs

To detect CpG DNA methylation at single base resolution, we performed WGBS on two replicates of pooled eggs, which gave us a six-fold coverage of the entire genome. The conversion rate of cytosines to uracil by the bisulfite treatment was 99.59 % and 99.58 % for the mapped reads for replicate one and two, respectively. For sites containing at least one unconverted cytosine the mean level of non-conversion was 0.3 %. Only 1038 partially unconverted CpGs occurred at the same site in both replicates. Overall 0.1 % of the cytosines at these sites were unconverted. Only 84 (8.1 %) of the sites with unconverted cytosines that were observed in both replicates mapped on a chromosome, while the rest was derived from unmapped regions. We also observed a highly significantly negative correlation between the fold coverage and the proportion of unconverted cytosines observed at the same CpG site (rho=-0.925, n=1038, p<0.001, Figure 1). This implies that the cytosines that remain after the bisulfite treatment can be either attributed to incomplete conversion or sequencing errors and therefore no CpG DNA methylation is present in the embryos.

**Figure 1.**
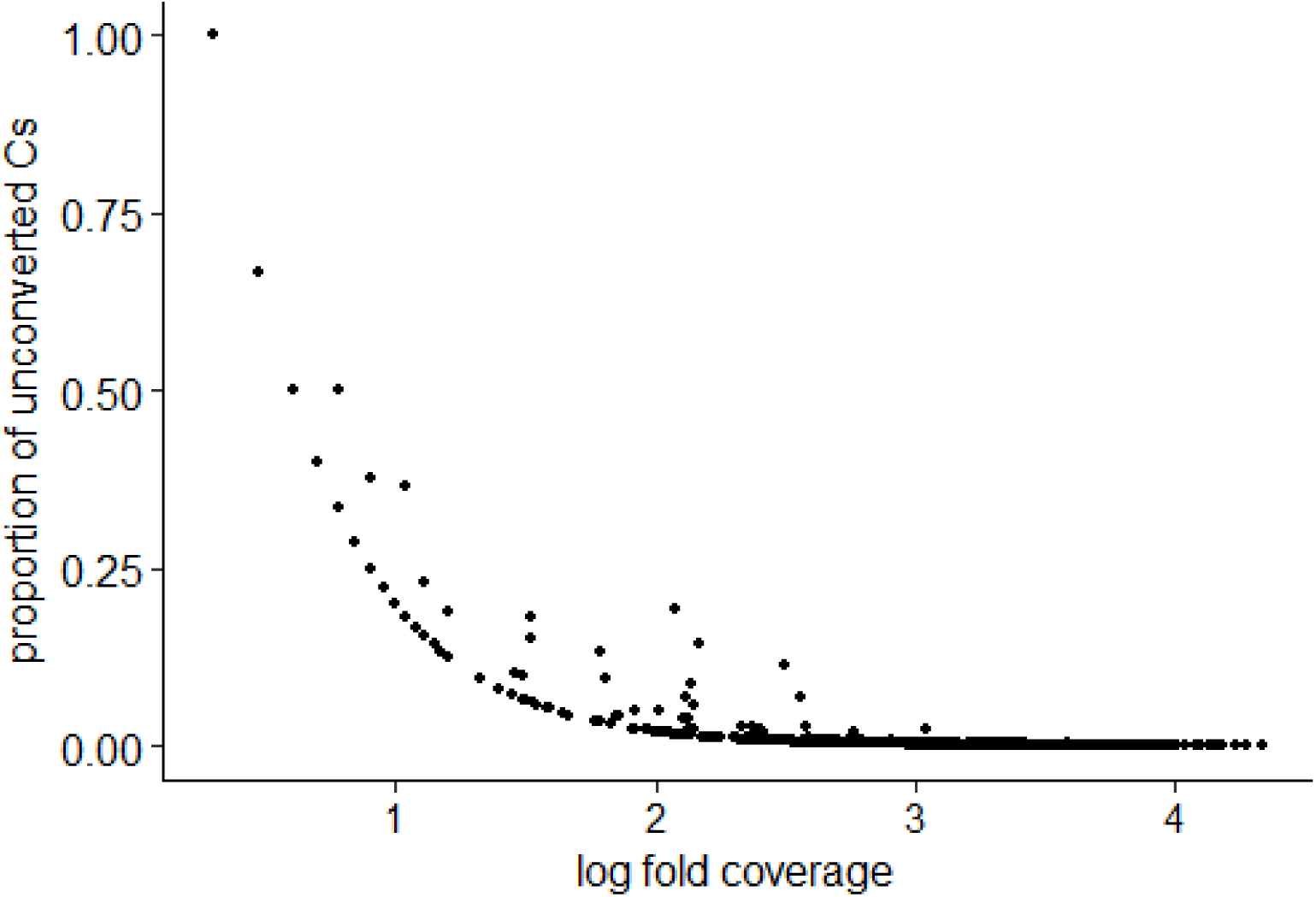
Correlation between the proportion of unconverted cytosines that were observed in both replicates of the WGBS and the log fold coverage at each site (Spearman rank, rho=-0.925, n=1038, p<0.001).

### 2.2 Expression of *Dnmt1*

To investigate the expression of *Dnmt1* across the entire life cycle of the beetle we analysed samples of eggs, larvae, pupae and adults via RT qPCR. We used equal numbers of male and female adults and pupae, while sex was not determined for eggs and larvae. All samples examined during this study contained measurable amounts of *Dnmt1* mRNA. The expression of *Dnmt1* was significantly higher in the eggs compared to whole body samples from adult beetles (*Table 1*). In larvae and pupae, we found *Dnmt1* to be expressed at similar levels as in the adult samples (*Table 1*). This suggested that *Dnmt1* might have a function throughout the whole life of the beetle.

**Table 1.** Gene expression of *Dnmt1* normalised over the expression of two housekeeping genes.

|  | relative expression<br>of *Dnmt1* | 95% C. I. | n | p value |
| --- | --- | --- | --- | --- |
| **Life stages (relative to adults)** |  |  |  |  |
| eggs | **1.660** | 1.027 - 3.144 | 4 | **0.004** |
| larvae | 0.818 | 0.511 - 1.555 | 8 | 0.063 |
| pupae | 0.794 | 0.390 - 1.784 | 8 | 0.176 |
| adults | 1.000 | 0.535 - 1.878 | 8 | 0.997 |
| **Gonads (relative to whole-body samples)** |  |  |  |  |
| male | 1.382 | 1.008 - 1.828 | 4 | 0.056 |
| female | **1.841** | 1.434 - 2.226 | 4 | **0.018** |
| **RNAi (relative to treatment control)** |  |  |  |  |
| **Females** |  |  |  |  |
| *Dnmt1* construct a | **0.680** | 0.301 - 1.226 | 11 | **0.002** |
| naive | 1.012 | 0.543 - 1.883 | 12 | 0.904 |
| **Eggs** |  |  |  |  |
| *Dnmt1* construct a | **0.442** | 0.226 - 0.753 | 3 | **0.025** |
| naive | 1.062 | 0.458 - 2.062 | 3 | 0.869 |
| **Females** |  |  |  |  |
| *Dnmt1* construct a | **0.646** | 0.553 - 0.752 | 4 | **0.011** |
| *Dnmt1* construct b | **0.703** | 0.536 - 0.906 | 4 | **0.021** |
| naive | 1.007 | 0.865 - 1.199 | 4 | 0.856 |
| **Males** |  |  |  |  |
| *Dnmt1* construct a | **0.686** | 0.542 - 0.822 | 3 | **<0.001** |
| naive | 1.062 | 0.853 - 1.312 | 3 | 0.458 |

We dissected virgin beetles of both sexes to investigate the expression of *Dnmt1* in the gonads. Any involvement in transgenerational processes would probably require the expression of the gene in the reproductive tissues so that mRNAs could be passed on to the offspring. In the female samples *Dnmt1* was significantly upregulated in the ovaries, while there was no significant difference in the expression of *Dnmt1* between the testes and whole-body samples in the males (*Table 1*).

### 2.3 Knockdown via maternal RNAi affects development and fecundity

To investigate possible functions of *Dnmt1* a knockdown via maternal RNAi was established. Firstly, we confirmed that the injection of *Dnmt1* dsRNA into female pupae led to a significant down-regulation in comparison to RNAi control injections, as measured by RT qPCR in the adult beetles. The treatment with *Dnmt1* dsRNA significantly reduced the expression of the target gene compared to the RNAi control, while expression did not differ significantly between the untreated and RNAi control (Table *1*). The knockdown was also transferred to the offspring, as the eggs of treated mothers showed significantly reduced amounts of *Dnmt1* dsRNA compared to eggs from the RNAi control (Table *1*).

Furthermore, *Dnmt1* knockdown significantly reduced the eclosion rate from pupae to imago (GLMM with binomial data distribution, df=2, X^2^ =146.2, p<0.001), suggesting that the protein plays a role in metamorphosis. Following *Dnmt1* knockdown, the eclosion rate three days post injection was 80 %, which was significantly lower than for the RNAi control (90 %) (Tukey’s HSD: z=6.93, p<0.001; Figure 2a). The naïve control achieved 96 % survival, which is significantly better than the RNAi control (Tukey’s HSD: z=-4.74, p<0.001; Figure 2a). This can be explained by the absence of injection wounds in these animals.

**Figure 2.**
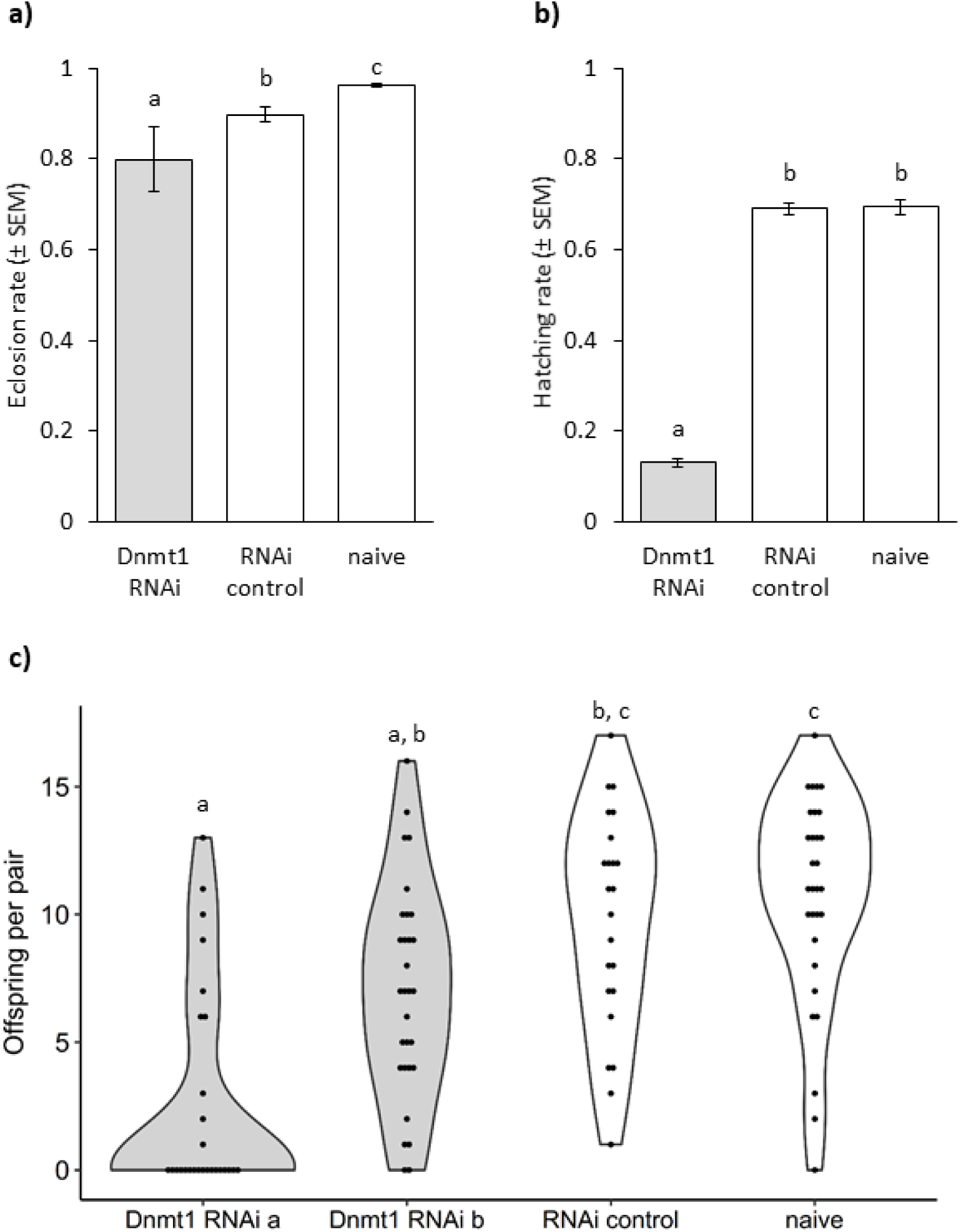
Phenotypic effects of maternal RNAi. **a)** Eclosion and survival rate (± SEM) of female pupae after *Dnmt1* RNAi treatment. Three replicates with 240-270 individuals per treatment and block. **b)** Larval hatching rates (±SEM) after mothers received *Dnmt1* RNAi treatment. Three replicates with 100 individualized eggs per treatment and block. **c)** Offspring produced by single mating pairs (n=26-30) after maternal RNAi treatment (*Dnmt1* RNAi construct a or b, RNAi or naïve control). Letters indicate significant differences.

We determined the fecundity of the treated beetles, by calculating hatching rates for one hundred eggs per treatment and experimental replicate. This showed that the fitness of the females, which had received the *Dnmt1* knockdown treatment was severely affected, as the hatching rate of their offspring was significantly reduced and only 13 % of larvae hatched, compared to 69 % in both controls (GLMM with binomial data distribution, df=2, X^2^=274.46, p<0.001; Tukey HSD: *Dnmt1* − RNAi control z=12.7, p<0.001, *Dnmt1* − naïve z=12.8, p<0.001, Figure 2b).

To confirm that the observed reduced fecundity was produced by the knockdown of *Dnmt1* and not any potential off-target effect on a different gene, we used a second non-overlapping construct. This construct also caused a significant downregulation in adults (Table 1) and significantly reduced female fecundity (Kruskal-Wallis df=3, X^2^=40.7, p<0.001, Figure 2c) compared to the naïve (Construct a: W=731.5, p<0.001, Construct b: W=675.5, p=0.001) and RNAi control (Construct a: W=553.5, p<0.001, Construct b: W=470, p=0.03). The two controls did not differ significantly from each other (W=301, p=0.44).

Additionally, we also investigated the effect of *Dnmt1* expression on male fertility. A knockdown in male pupae significantly reduced *Dnmt1* expression in the adults (Table *1*) but did not reduce their fertility over the first nine days of mating compared to the RNAi control or naïve group (GLMM with negative binomial data distribution, df=2, X^2^=2.3441, p-value=0.31).

### 2.4 *Dnmt1* knockdown disrupts embryonic development

We also performed a closer analysis of embryos after maternal knockdown of *Dnmt1.* After twelve days, hardly any larvae had hatched, and we therefore assumed that the eggs had either not been fertilised or that the knockdown was causing an embryonic developmental arrest, which was previously observed in *Nasonia vitripennis* ^13^. In order to identify the time point of a potential developmental arrest, we sampled eggs every two hours for the first ten hours after oviposition. According to their developmental status, the eggs were then assigned to one of four categories of a timeline of normal embryonic development. At the two first time points, we did not observe any significant developmental differences between the treatment groups (age 0-2 h: OLM _2, 96_,χ2=0.84, p=n.s.; age 2-4 h: OLM _2, 140_, χ2=4.65, p=0.098). But, when the embryos were between four and six hours old, more than 60 % of the eggs of the knockdown treatment fell within the earliest category of development and therefore differ significantly from the two controls (OLM_107_, χ2=35.71, p<0.001). We observed a similar pattern at the embryonic ages of six to eight hours (OLM_105_, χ2=55.11, p<0.001) and eight to ten hours (OLM_144_, χ2=40.76, p<0.001). We therefore conclude that the knockdown of *Dnmt1* causes an early embryonic arrest after first few cleavage cycles (Figure 3).

**Figure 3.**
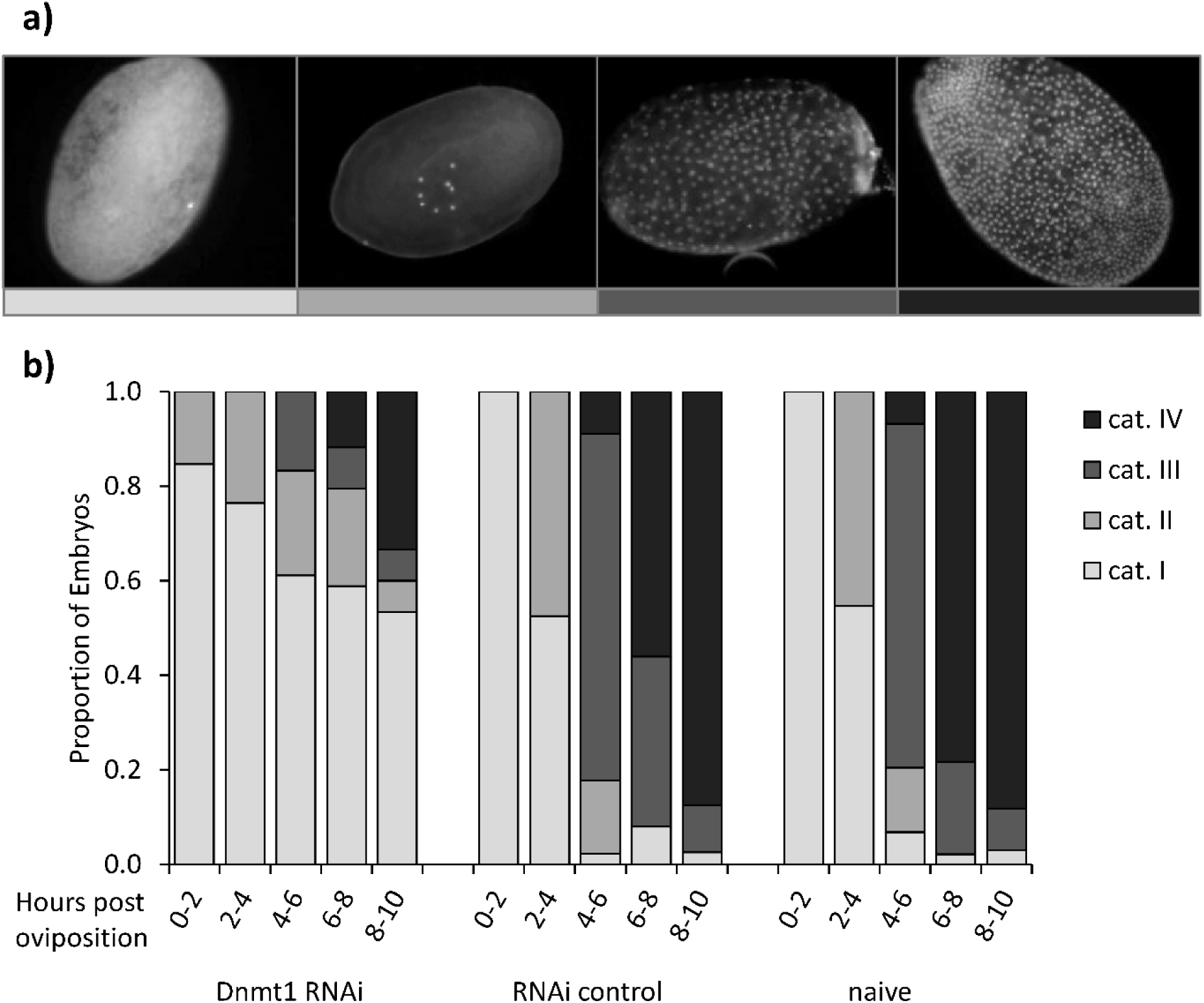
Dnmt1 knockdown and embryonic development. **a)** Timeline of naïve embryonic development. Shown is (left to right) a representative example for each of the following age categories: cat. I (ca. 1h), cat. II (ca. 3h), cat. III (ca. 5h), cat. IV (ca. 8h) after oviposition. Eggs were produced by untreated animals. **b)** Phenotypes of embryos after maternal RNAi. Eggs from the treatment and controls (*Dnmt1* RNAi, RNAi control and naïve) and five age groups (hours post oviposition) were sorted according to their developmental status into one of four categories established from the timeline of naïve development.

## 3 Discussion

Using WGBS, we did not observe any recognizable evidence for CpG DNA methylation in *T. castaneum* embryos. The few candidate methylation marks detected by whole genome sequencing are most likely due to incomplete bisulfite conversion or sequencing errors. Furthermore, there is hardly any overlap in the candidate mCpGs between the two replicates, which leads us to conclude that CpG DNA methylation is absent in two-day old embryos. Our results agree with those of Zemach *et al.,* who were also unable to detect any significant levels of CpG DNA methylation in *T. castaneum* ^19^. Yet, two other studies found DNA methylation in all life stages of the beetle. While Song *et al.* (2017) observed a strong preference for CpA methylation, which because of its asymmetrical nature is very unlikely to be produced by DNMT1, Feliciello *et al.* (2013) found CpG methylation especially on satellite DNA but also spread across the genome of embryos. These differences might be explained by changing methylation patterns during different life stages and possible rapid dynamic changes during embryonic development similar to the epigenetic reprogramming in mammals ^34^. Alternatively, they might also result from incomplete bisulfite deamination, which is often linked to the artefactual detection of methylation outside of the canonical CpG context.

*T. castaneum,* as several other insect species, lacks *Dnmt3* in its genome and therefore does not possess a complete DNA methylation system ^3,29,43^. Nevertheless, some insect species lacking a complete set of *Dnmts,* show detectable DNA methylation including diverse methylation patterns in correlation with environmental changes ^15,16,44^. It is still unknown whether *Dnmt1* can take over the function of the missing *Dnmt3* or if other enzymes are responsible for preserving the function. However, the lack of CpG methylation observed in the present study is further supported by the absence of CpG depletion from the beetle’s gene set, which is a known evolutionary signal of CpG methylation ^32,45^. To ultimately resolve this controversy, further studies are needed to determine whether, where and at which time functional levels of CpG DNA methylation might occur in *T. castaneum.*

Our results also show that *Dnmt1* is present and expressed in *T. castaneum* during all life stages, in both sexes and in the reproductive tissues despite the apparent absence of CpG DNA methylation. The expression of *Dnmt1* during the entire life cycle of the beetle hints at a possible function during several life stages. Because ovaries and eggs showed significantly higher relative amounts of *Dnmt1* mRNA, the protein might be especially relevant during reproduction and embryonic development.

Both constructs of *Dnmt1* used in our RNAi experiment significantly reduced the levels of *Dnmt1* mRNA. Although the reduction was only to 70% compared to the levels observed in the treatment control, it still produced a strong phenotype in its effect on offspring embryonic development. RNAi treatment in *B. mori* leads to a similar degree of reduction of *Dnmt1* and also decreased hatchability ^44^. A recently described alternatively spliced isoform of *Dnmt1* in *T. castaneum* lacks the C-terminal DNA methylase domain and is thus unable to methylate DNA, but was shown to be predominantly expressed ^36^. While the dsRNA constructs used here should cause a knockdown of both isoforms, our gene expression analysis was only able to detect the transcript containing the methylase domain. Thus, the RNAi treatment might affect the predominant shorter isoform to a greater extent.

We could show that *Dnmt1* is essential for early embryonic development even in the absence of functional levels of CpG DNA methylation. The knockdown in mothers leads to high lethality in the offspring embryos within the first hours after oviposition. Similarly, the maternal knockdown of one copy of *Dnmt1* in *N. vitripennis* also lead to a developmental arrest in the embryos but at a later time point ^13^. Additionally, fecundity in brown planthoppers was strongly decreased after *Dnmt1* knockdown ^46^. DNMT1 appears to also have an essential role in metamorphosis of *T. castaneum* as a reduction during the pupal stage decreased the rate of eclosion. It is therefore reasonable to assume that DNMT1 has additional functions in *T. castaneum* and maybe also in other insects, including those possessing a DNA methylation machinery.

Alternative, non-catalytic functions of *Dnmt1* have been suggested ^47^. The DNA-binding function could be intact in the absence of methylase domain, still enabling the enzyme to fulfil other tasks ^36^. This is supported by the presence of a large N-terminal domain without methylase function but with many molecular binding patterns ^47^. For example, specific motifs of the N-terminal domain have been shown to regulate the transcription of E-cadherin in humans without the association with any changes in methylation patterns ^48^. Further studies in vertebrates indicate that DNMT1 binding to histone modification enzymes is involved in modulation of gene expression ^49,50^. Additional support comes from the clawed frog *Xenopus laevis,* for which *Dnmt1* expression is also essential during embryonic development ^51^. Here, the protein silences genes by directly repressing transcription independent of its catalytic function ^52^. The downregulation of *Dnmt1* leads to the premature activation of gene expression before the mid-blastula transition without any larger changes in methylation patterns ^52^. This may indicate that also in insects including *T. castaneum* the sufficient provision of maternal *Dnmt1* transcripts ensures the correct timing of the maternal to zygotic transition of mRNA transcription and thereby prevents embryonic developmental arrest and death. The results of this study emphasise a presumably non-catalytic but essential role of *Dnmt1,* which needs to be investigated in more detail.

## 4 Methods

### 4.1 Model organism

We used beetles from the Georgia 2 (GA-2) line for WGBS, since the genome sequence of *T. castaneum* was produced from that line ^29^. Expression analysis and knockdown experiments were carried out with individuals from the San Bernadino (SB) line originating in the Sokoloff laboratory (Tribolium Stock Center, California State University, San Bernardino, USA). Beetles were kept on organic wheat flour (type 550) with 5 % brewer’s yeast at a population size of more than 200 individuals. They were maintained at 30 °C and 70 % humidity at a 12h-light/12h-dark cycle.

### 4.2 Whole genome bisulfite sequencing

WGBS was performed on two replicates each from 300-400 μl of eggs. We collected eggs after a 24 hours oviposition period from a population of about 1000 individuals of the GA-2 strain. Samples were incubated at 30 °C with 70 % humidity for an additional 24 h, then frozen in liquid nitrogen, and stored at −80 °C.

Per sample 5 μg of high molecular weight DNA were used for fragmentation using the Covaris S2 AFA System in a total volume of 100 μl. The fragmentation (parameters: duty cycle: 10 %; intensity: 5; cycles/burst: 200; cycles: 3) was run for a total fragmentation time of 180 s. Fragmentation was confirmed with a 2100 Bioanalyzer (Agilent Technologies) using a DNA1000 chip. The fragmented DNAs were concentrated to a final volume of 75 μl using a DNA Speed Vac. End repair of fragmented DNA was carried out in a total volume of 100 μl using the Paired End DNA Sample Prep Kit (Illumina) as recommended by the manufacturer. For the ligation of the adaptors, the Illumina Early Access Methylation Adaptor Oligo Kit and the Paired End DNA Sample Prep Kit (Illumina) were used, as recommended by the manufacturer. For the size selection of the adaptor-ligated fragments, we used the E-Gel Electrophoresis System (Invitrogen) and a Size Select 2 % precast agarose gel (Invitrogen). Each fragmented DNA was loaded on two lanes of the E-gel. Electrophoresis was carried out using the “Size Select” program for 16 min. According to the standard loaded (50 bp DNA Ladder, Invitrogen), 240 bp fragments were extracted from the gel, pooled, and directly transferred to bisulfite treatment without further purification. For the bisulfite treatment we used the EZ-DNA Methylation Kit (Zymo) as recommended by the manufacturer with the exception of a modified thermal profile for the bisulfite conversion reaction. The conversion was carried out in a thermal cycler using the following thermal profile: 95 °C for 15 s, 50 °C for 1 h, both steps repeated 15 times, 4 °C for at least 10 min. The libraries were subsequently amplified, using the Fast Start High Fidelity PCR System (Roche) with buffer 2, and PE1.1 and PE2.1 amplification primers (Illumina). PCR thermal profile: 95 °C for 2 min, 95 °C for 30 s, 65 °C for 20 s, 72 °C for 30 s, then repeat from step 2, 11×, 72 °C for 7 min, hold at 4 °C. PCR reactions were purified on PCR purification columns (MinElute, Qiagen) and eluted in 20 μl elution buffer (Qiagen).

Paired-end sequencing was performed on an Illumina HiSeq 2000 system with read lengths of 105 bp and an average insert size of 240 bp. Reads were trimmed and mapped with BSMAP 2.5 ^53^ using the assembly version 3.0 of the *T. castaneum* genome (https://www.hgsc.bcm.edu) as a reference sequence. Duplicates were removed using the Picard tool (http://broadinstitute.github.io/picard). Methylation ratios were determined using a Python script (methratio.py) distributed together with the BSMAP package.

For every site containing a CpG, we calculated the fold coverage and the proportion of cytosines at each site that had not been converted by the bisulfite treatment. We also computed which cytosines that were partially unconverted occurred in both replicates and combined this data. Finally, we checked whether there is a correlation between proportion of unconverted cytosines and fold coverage.

### 4.3 Gene Expression Analysis

We measured the expression of *Dnmt1* in different life history stages (eggs, larvae, pupae, adults) as well as in the reproductive organs of the beetle using quantitative real-time PCR (RT qPCR). At the time point of collection, around 200 μl of eggs were used per sample, while for all other samples material from five individuals was pooled, immediately shock frozen in liquid nitrogen and stored at −80 °C. We analysed four replicates for each life history stage as well as for the gonad and whole-body samples.

The applied RNA extraction protocol combines phenol/chloroform lysis and extraction with the purification via spin columns from the SV Total RNA Isolation System (Promega) and was performed as described in Eggert et al. (2014). We performed reverse transcription into complementary DNA (cDNA) with the RNA-dependent DNA polymerase SuperScript III and the use of oligo(dT)18 primers (Invitrogen). The reaction started with 100 ng RNA and produced a final volume of 10 μl. The samples were stored at 4 °C for up to one week. For all samples, we measured the expression of *Dnmt1* relative to the expression of two housekeeping genes the *ribosomal protein L13a* (*RpL13a*) and *ribosomal protein 49* (*rp49*). All primers were designed to cross exon-intron boundaries and their quantification efficiency (E) was calculated using a serial dilution curve (Supplementary Table S1).

We applied qPCR using the LightCycler^®^ 480 Real-Time PCR System (Roche) and KAPA SYBR^®^ FAST qPCR Light Cycler 480 reaction mix (PEQLAB). Prior to the run of the assay, we diluted the cDNA 1:10. All experiments were carried out with two technical replicates per sample. Crossing points (Cp) for each technical replicate were calculated and used in further analysis if the standard deviation (STD) between the two technical replicates was below 0.5.

### 4.4 Knockdown via parental RNAi

To accomplish a knockdown of *Dnmt1* via parental RNAi, we designed a construct for double stranded RNA (dsRNA). DsRNA complimentary to the *Asparagin synthetase A* gene (*AsnA*) from *Escherichia coli* served as a treatment control ^55^. We produced both dsRNA constructs using EcoRV digested pZErO^TM^-2 vector (Invitrogen) and *in vitro* transcribed into dsRNA with the T7 RNA polymerase based T7 MEGAscript Kit (Ambion). iBeetle-Base provided us with the sequence for the second non-overlapping construct, which we used in order to be able to exclude possible off-target effects (Schmitt-Engel et al. 2015, http://ibeetle-base.uni-goettingen.de/details/iB_08496). For this we obtained the dsRNA from EupheriaBiotech.

For the RNAi procedure, we treated female and male pupae with either *Dnmt1* dsRNA, *AsnA* dsRNA or left them naïve, *i.e.* untreated. For the injections, we adjusted the concentration of dsRNA to 1,800 ng/μl. Prior to the injections, three-day old pupae were collected and glued with the end of their abdomen to microscopic slides with Fixogum (Marabu). The injections were carried out with a nano injector (FemtoJet, Eppendorf) using borosilicate glass capillaries (100 mm length, 1.0 mm outside diameter, 0.021 mm wall thickness; Hilgenberg). Pupae were injected with the dsRNA solution laterally between the second and third segment of the abdomen. We stopped the injections when the pupae stretched, due to the increasing turgor.

The pupae were left on the glass slides in Petri dishes until they eclosed one to five days later. After eclosion individual survival was recorded and the surviving beetles could mature for four days. At this point, we randomly sampled individuals for either expression analysis or mating and egg production. Due to large sample sizes required in the experiment involving embryo staining, we conducted these injections in three consecutive experimental blocks. We confirmed the success of the knockdown by performing expression analysis on four pools of three to five individuals per treatment and block. Additionally, we measured the *Dnmt1* mRNA amount present in the eggs, by performing RT qPCR on one pool of 150-200 μl eggs per each of the three experimental blocks and treatment groups. At the time point of RNA extraction eggs of each sample were between 45 minutes and 16 hours old.

### 4.5 Fitness costs of the knockdown

To estimate potential fitness costs of the *Dnmt1* knockdown treatment we collected one hundred eggs from each of the treatment groups and controls, individualised them and counted live larvae twelve days later. For the second non-overlapping dsRNA construct (construct b) fitness costs were examined in a different way. The construct from the previous experiment (construct a) was also used here. We set up single mating pairs (n=23-30) and let them mate for 24 hours. Then live larvae were counted twelve days later for each pair. Besides the two dsRNA constructs, a RNAi control and a naïve group were included.

To measure the fertility of the males after RNAi treatment, individual mating pairs with untreated, virgin females of the same age were formed four days post eclosion. The pairs were put on to new flour after 24 hours and then again, every three days for the next nine days. Two weeks after the oviposition living larvae were counted for each pair (n=20-26).

### 4.6 Embryonic development after maternal knockdown

To determine a possible effect of the maternal knockdown on their offspring, we observed the embryonic development via DNA staining. For this we collected eggs every two hours over a period of ten hours, which enabled us to look at five individual time points after oviposition. For the fixation, we washed the eggs with water and treated them with 25 % DanKlorix (Colgate –Palmolive), which contains 5% sodium hypochlorite to remove the chorion. The eggs were submerged in heptane and fixed with 4 % formaldehyde in phosphate buffered saline (PBS). Vigorous shaking in methanol led to the removal of the vitelline membrane. Finally, the methanol was replaced with 95% ethanol and the fixed embryos were stored at −20 °C.

We determined the developmental stage of the previously fixed embryos through staining with 4’,6-diamidino-2-phenylindole (DAPI) (Carl Roth). After incubating the embryos in PBS with 0.5 μg/ml DAPI for five minutes, they were washed with PBS twice and mounted to a microscopy glass slide using Fluoroshield histology mounting medium (Sigma-Aldrich). A fluorescence inversion microscope (Observer Z1, Zeiss) with an attached fluorescence camera (AxioCamMR3, Zeiss) was used for taking pictures at 200x magnification. We ascertained embryonic development by assigning each picture to one of four categories. These categories were formed from a time line of naïve development over the first ten hours after oviposition (cat.I=ca. 1h, cat.II=ca. 3h, cat.III=ca. 5h, cat.IV=ca. 8h), which was established prior to this experiment.

### 4.7 Statistical analysis

If not stated otherwise, all statistical analyses were performed in R (R Development Core Team 2008, version 3.4.0) under RStudio version 0.99.467 ^58^ using packages lme4 ^59^ and MASS ^60^ when performing generalised linear mixed effects models (GLMMs). The correlation between fold coverage and proportion of unconverted cytosines was tested using a Spearman rank test.

All expression data of *Dnmt1* were analysed and tested for significant differences using the relative expression software tool REST2009 (Qiagen ; Pfaffl, Horgan, & Dempfle, 2002) as previously described ^54^. Bonferroni correction for multiple testing was applied to the results if the control group was compared to more than one treatment group.

For examining the eclosion and therefore survival after injection of the females in the knockdown experiments a GLMM with binomial error distribution was applied to the data. We investigated the effect of the knockdown on maternal fitness by testing hatching rates of larvae for differences in a GLMM with binomial error distribution followed by a Tukey’s HSD test. Fecundity of individual pairs after maternal knockdown using both *Dnmt1* RNAi constructs was tested with a Kruskal-Wallis test, then pairwise comparisons were performed with a Wilcoxon test and p values were adjusted according to Benjamini & Hochberg (1995). Male fertility after knockdown was analysed with a generalised linear model (GLM) with negative binomial error distribution.

The embryonic development after maternal knockdown was tested with an ordinal logistic model (OLM) and a post-hoc maximum likelihood test to determine, whether the distribution of embryos to the four developmental categories was significantly different between the maternal treatments. This was done for each observed time point separately. Analyses were performed with JMP version 8 (SAS Institute Inc).

## 5 Data availability

The datasets generated during and/or analysed during the current study are available from the corresponding author on reasonable request.

## 7 Acknowledgement

This work was supported in part by the Volkswagen Stiftung, project number I/84 794.

## 8 Author contributions

Conceived and designed study: NS, IW, MDdB, FL, JK. Carried out lab work: NS, IW, JE. Carried out sequencing: GR. Analysed data: NS, GR. Wrote manuscript: NS with comments from all other authors.

